# Focal radiotherapy improves CAR T cell therapy targeting prostate cancer

**DOI:** 10.64898/2026.06.23.734073

**Authors:** Cari A. Young, Jiangyue Liu, Yuwei Ren, Reginaldo Rosa, Lea Christian, Handan Hong, Lupita Lopez, Alyssa Buckley, Jingting Hao, Yukiko Yamaguchi, Anthony K. Park, Hemendra Ghimire, Amr Mohamed Hamed Abdelhamid, Darren Zuro, Susanta Hui, Catalina Martinez, Stephen J. Forman, Yun Rose Li, Tanya B. Dorff, John P. Murad, Saul J. Priceman

## Abstract

Chimeric antigen receptor (CAR) T cell therapy has limited efficacy against solid tumors such as prostate cancer due to the immunosuppressive tumor microenvironment (TME). Combining CAR T cells with existing therapies that remodel the TME and promote endogenous immune responses, such as radiation therapy and chemotherapies, may strengthen antitumor responses. Here, we assessed the potency of combining focal radiotherapy (RT), cyclophosphamide (Cy) preconditioning, and prostate stem cell antigen (PSCA)-CAR T cells against syngeneic prostate cancer models. Focal RT alone increased T cell and dendritic cell infiltration and activation in the irradiated tumor. Furthermore, the combination of all three therapies was critical for enhanced antitumor responses and survival across multiple subcutaneous, bone-metastatic, and multifocal disease models. This combination, in the irradiated TME and tumor-draining lymph nodes (tdLN), led to greater antigen presentation by myeloid cells and endogenous T cell activation and cytotoxicity. Our study demonstrates the potency of combining focal RT with PSCA-CAR T cells, significantly improving therapeutic responses in the irradiated tumor and contributing to a more robust systemic immune response against metastatic burden in prostate cancer.

## Introduction

Chimeric antigen receptor (CAR) T cell therapies have demonstrated clinical successes against hematological malignancies; however, the development of successful CAR T cells against solid tumors, particularly immunologically “cold” tumors like metastatic castration-resistant prostate cancer (mCRPC), is challenging due to the immunosuppressive tumor microenvironment (TME)[1, 2]. Combining CAR T cells with other therapies that can remodel the TME and increase immune infiltration prior to CAR T cell infusion can help promote stronger therapeutic responses. CAR T cells directed against the prostate stem cell antigen (PSCA) in solid tumors require preconditioning with lymphodepleting agent cyclophosphamide (Cy), which promotes durable antitumor responses, greater T cell infiltration, and pro-inflammatory signals within the prostate TME[3]. In our recent phase 1 clinical trial evaluating PSCA-CAR T cells therapy in patients with mCRPC, we found greater expansion of peripheral blood CAR T cells and therapeutic responses with lymphodepletion prior to CAR T cell infusion[4]. However, the magnitude and durability of the responses in both preclinical and clinical settings have been limited, suggesting that additional T cell engineering and/or combinatorial strategies are needed.

Radiation therapy (RT) is a standard of care for cancer and is used in both the curative and palliative settings [5, 6]. Beyond the cytotoxic effects on tumor cells, RT has numerous immunomodulatory effects in the TME, including enhancing antigen presentation and inducing immunogenic cell death, which activates innate and adaptive immunity[7, 8]. These immunostimulatory effects suggest that RT may increase immunotherapy response rates. Clinical trials combining RT with immune checkpoint inhibitors have demonstrated improved survival compared to treatment with immune checkpoint inhibitors alone against non-small cell lung carcinoma, melanoma, and mCRPC[9–13]. Preclinical studies combining RT with CAR T cells have also demonstrated more effective CAR T cell killing and tumor control with prior irradiation of the tumor in mesothelioma, triple-negative breast cancer, and pancreatic cancer models[14–16]. These studies suggest that the combination of focal RT and CAR T cell therapy may also effectively treat prostate cancer, leading to more durable antitumor efficacy and improvements in overall survival.

To this end, we examined the effects of combining focal RT with PSCA-CAR T cells in multiple models of prostate cancer. Using heterozygous human PSCA-knockin (hPSCA-KI) immunocompetent mouse models that express both hPSCA and murine PSCA in normal tissues, we found that focal RT alone inhibited prostate tumor growth and induced immune cell infiltration within the irradiated TME. We then assessed the antitumor response of combining RT, Cy lymphodepletion, and PSCA-CAR T cells against subcutaneous tumors and intratibial prostate tumor models representing a bone-metastatic setting and found in both models that this triple combination significantly improved antitumor responses and survival compared to either combination alone. Next, we characterized the effects of varying radiation doses and fractionation in a multifocal disease model and identified a dose that induced the greatest antigen presentation in the locally irradiated TME. Using the same multifocal disease model, we found that this combination of treatments consistently led to the greatest survival benefit. Mechanistically, we found that this combination treatment led to higher levels of T cell activation and upregulation of antigen presentation markers in the TME and in tumor-draining lymph nodes.

These studies demonstrate a translatable combination strategy to improve the efficacy of PSCA-CAR T cells against mCRPC and are informative for ongoing clinical trials.

## Results

### Focal radiotherapy modulates the immune landscape in the prostate TME

We first aimed to assess the effects of focal RT treatment on the growth and local TME of mouse prostate tumors. Prior studies found that 16 Gy total radiation given in 2 fractions significantly inhibited the growth of the Ras- and Myc-transformed mouse prostate carcinoma cell line RM-1 and induced changes in T cell and myeloid cell infiltration over time[17]. To verify these effects, we engrafted the related mouse prostate tumor cell line, RM-9, engineered to express the human PSCA antigen (RM9-hPSCA). The tumors were irradiated with 16 Gy total radiation given in two fractions (16Gy in 2) (**Figure 1a**). This dose of radiation significantly inhibited tumor growth (**Figure 1b**), as observed previously with the RM-1 model.

**Figure 1.**
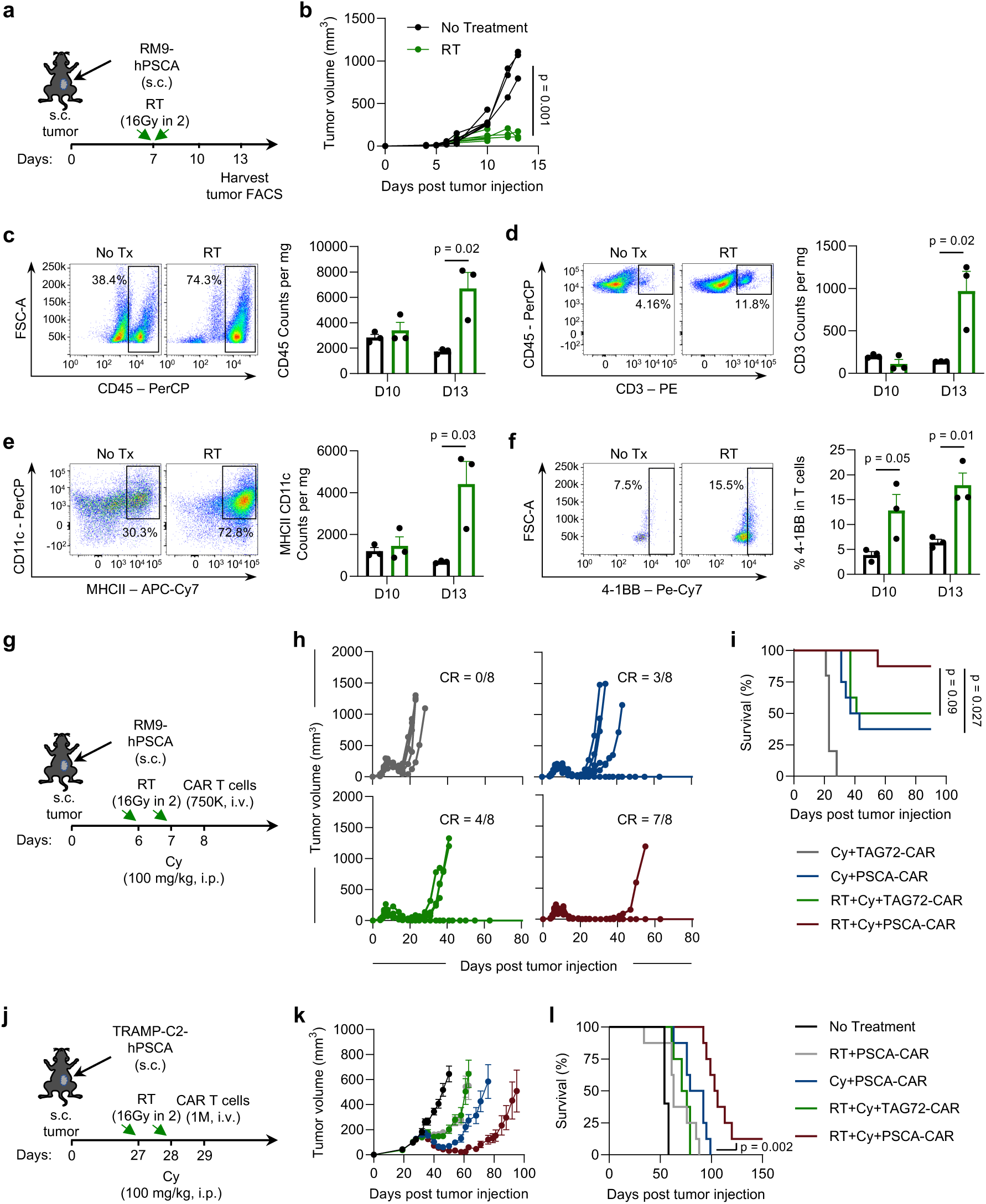
Focal RT modulates the immune landscape of the prostate TME and improves survival when combined with Cy preconditioning and PSCA-CAR T cells. **a**, Illustration of RM9-hPSCA s.c. tumor engraftment and treatment with focal RT (16 Gy total in 2 fractions given 6 hours apart) followed by tumor collection and flow cytometric analysis on the indicated days. **b**, Tumor volume (mm^3^) measurements of each replicate at indicated days post tumor injection for indicated treatment groups. **c-f**, Representative flow cytometry plots of samples harvested 13 days after tumor engraftment (or 6 days after RT) showing percentages and counts per mg of CD45+ immune cells (**c**), counts per mg of CD45+ CD3+ T cells (**d**), counts per mg of CD11c+MHCII+ antigen-presenting dendritic cells (**e**) and percentage of 4-1BB+ activated T cells (**f**) in tumors harvested 10 and 13 days after engraftment, or 3 and 6 days after RT treatment, respectively. **g**, Illustration of RM9-hPSCA s.c. tumor engraftment and treatment with focal RT (16 Gy total in 2 fractions given 24 hours apart), Cy preconditioning (100 mg/kg, i.p. following the second RT dose), and 0.75 x 10^6^ T cells (TAG72-CAR or PSCA-CAR, i.v.) on the indicated days. **h**, Tumor volume (mm^3^) measurements of each replicate at indicated days post tumor injection for treatment groups; CR, complete response. **i**, Kaplan-Meier survival plot for mice in each indicated group; n=8 mice per group, p-value compares RT+Cy+PSCA-CAR and Cy+PSCA-CAR or RT+Cy+TAG72-CAR using a log-rank (Mantel-Cox) test. **j**, Illustration of TRAMPC2-hPSCA s.c. tumor engraftment and treatment with focal RT (16 Gy total in 2 fractions given 24 hours apart), Cy preconditioning (100 mg/kg, i.p. following the second RT dose), and 1 x 10^6^ T cells (TAG72-CAR or PSCA-CAR, i.v.) on the indicated days. **k**, Average tumor volumes presented as mean ± SEM at indicated days post tumor injection for treatment groups. **l**, Kaplan-Meier survival plot for mice in each indicated group; n=8 mice per group, p-value compares RT+Cy+PSCA-CAR and Cy+PSCA-CAR using a log-rank (Mantel-Cox) test.

To investigate the immunological changes in the local TME induced by RT, we harvested the tumors three and six days following RT treatment, corresponding to day 10 and 13 post tumor engraftment, respectively, for flow cytometric analysis. Although minimal changes in immune cell infiltration were observed three days following RT, there were significant increases in CD45+ immune cell and CD3+ T cell infiltration six days following RT (**Figure 1c-d**). We also observed higher infiltration of CD11c+MHC-II+ antigen-presenting dendritic cells (DCs) six days following RT (**Figure 1e**). Interestingly, 4-1BB in T cells was higher three days following RT, suggesting early T cell activation preceded changes in T cell infiltration (**Figure 1f**). These results indicate that 16Gy in 2 fractions induced immune-stimulating changes on the local TME.

### The combination of focal radiotherapy, Cy preconditioning, and PSCA CAR T cell treatments improves survival against multiple mouse prostate cancer models

Next, we wanted to assess the effects of focal RT on PSCA-CAR T cell efficacy in our mouse prostate cancer model. Previously our group showed that Cy preconditioning was necessary for enhancing PSCA-CAR T cell therapy against immunosuppressive mouse prostate and pancreatic tumors[3]. We hypothesized that focal RT alone given prior to CAR T cell treatment would have a similar beneficial effect and lead to curative responses. We treated subcutaneous (s.c.) RM9-hPSCA mouse prostate tumors with Cy or RT prior to PSCA-CAR T cell treatment (**Supplementary Figure 1a**). While Cy+PSCA-CAR T cell treatment cured 4 out of 6 mice, showing an antitumor response consistent with our previous studies, treatment with RT+PSCA-CAR T cells transiently inhibited tumor growth but failed to show curative responses (**Supplementary Figure 1b-c**). These results showed that RT alone does not sufficiently induce immunostimulatory effects on the local TME to improve PSCA-CAR T cell activity and that Cy preconditioning may be critical for CAR T cell responses.

Using a suboptimal CAR T cell dose and a non-targeting control CAR T cell targeting tumor-associated glycoprotein 72 (TAG72-CAR), we tested different combinations of Cy and CAR T cells with or without prior RT treatment (**Figure 1g**). While Cy+TAG72-CAR T cell treatment transiently inhibited tumor growth, the Cy+PSCA-CAR T cell and RT+Cy+TAG72-CAR T cell treatments both cured about half of the mice (**Figure 1h**). The combination of focal RT+Cy+PSCA-CAR T cell treatment led to the greatest survival benefit with significantly better survival compared to the Cy+PSCA-CAR T cell treatment, highlighting the contribution of RT. (**Figure 1i**). Interestingly, we observed potent therapeutic responses with RT+Cy+TAG72-CAR T cells, supporting this combination even in the absence of targeted CAR T cell therapy; however, treatment with prostate cancer- targeting PSCA-CAR T cells led to improved survival, demonstrating the importance of treating with a targeting CAR to maximize the therapeutic effect. Collectively, these results suggest that focal RT greatly enhances the antitumor activity of Cy-preconditioned PSCA-CAR T cell therapy.

We next assessed the impact of these combination treatments on endogenous long-term antitumor immunity. Cured mice from the previous study that remained tumor-free for about 60 days were rechallenged with antigen-negative RM9-WT tumors. While one mouse from each of the Cy+PSCA-CAR T cell and RT+Cy+TAG72-CAR T cell pre-treated groups rejected tumors, the RT+Cy+PSCA-CAR T cell treatment led to the highest frequency of rejected tumors and trended towards better survival (**Supplementary Figure 2a-c**). This suggests that the combination therapy may induce a stronger protective immune memory response independent of PSCA expression.

To confirm the efficacy of the combination treatment against a second syngeneic prostate cancer model, we engrafted s.c. TRAMPC2-hPSCA prostate tumors and treated mice with different combinations of focal RT, Cy, and PSCA-CAR T cells (**Figure 1j**). RT+PSCA-CAR T cells and RT+Cy+TAG72-CAR T cells did not demonstrate curative responses (**Figure 1k**). While Cy+PSCA-CAR T cell treatments in this model showed greater antitumor response than the previous two groups, the combination of RT+Cy+PSCA-CAR T cells showed the greatest inhibition of tumor growth, curing 1 out of 8 mice, and significantly prolonged survival compared to the other treatment groups (**Figure 1k-l**). These results validate our findings in the RM9-hPSCA prostate cancer model and indicate that the combination of focal RT, Cy lymphodepletion, and PSCA-CAR T cells are critical for enhancing PSCA-CAR T cell efficacy against prostate cancer. Furthermore, although there was only one cured mouse from the RT+Cy+PSCA-CAR T cell treatment, we engrafted antigen-negative TRAMPC2-WT tumors into the cured mouse and observed complete rejection, which is consistent with the strong protective immune memory seen in the RM9 model (**Supplementary Figure 2d-e**).

Because prostate cancer frequently metastasizes to the bone, we next determined the efficacy of this combination therapy against a clinically relevant model of prostate cancer bone metastasis[18]. We engrafted RM9-hPSCA tumors into the tibia, treated mice with different combinations of focal RT, Cy, and CAR T cells, and monitored tumor growth using bioluminescent imaging (**Figure 2a**). The RT+Cy+TAG72-CAR T cell and the Cy+PSCA-CAR T cell conditions showed similar levels of tumor growth inhibition, with curative responses in 2 out of 10 mice treated with Cy+PSCA-CAR T cells (**Figure 2b-d**). In contrast, the combination of RT+Cy+PSCA-CAR T cells cured 10 out of 11 treated mice, indicating that the focal RT treatment greatly improved the PSCA-CAR T cell killing efficacy and overall survival (**Figure 2b-d**). These results show that the combination of focal RT, Cy preconditioning, and PSCA-CAR T cell treatment consistently leads to durable therapeutic responses and improved survival in multiple models, including a clinically relevant bone-metastatic disease model.

**Figure 2.**
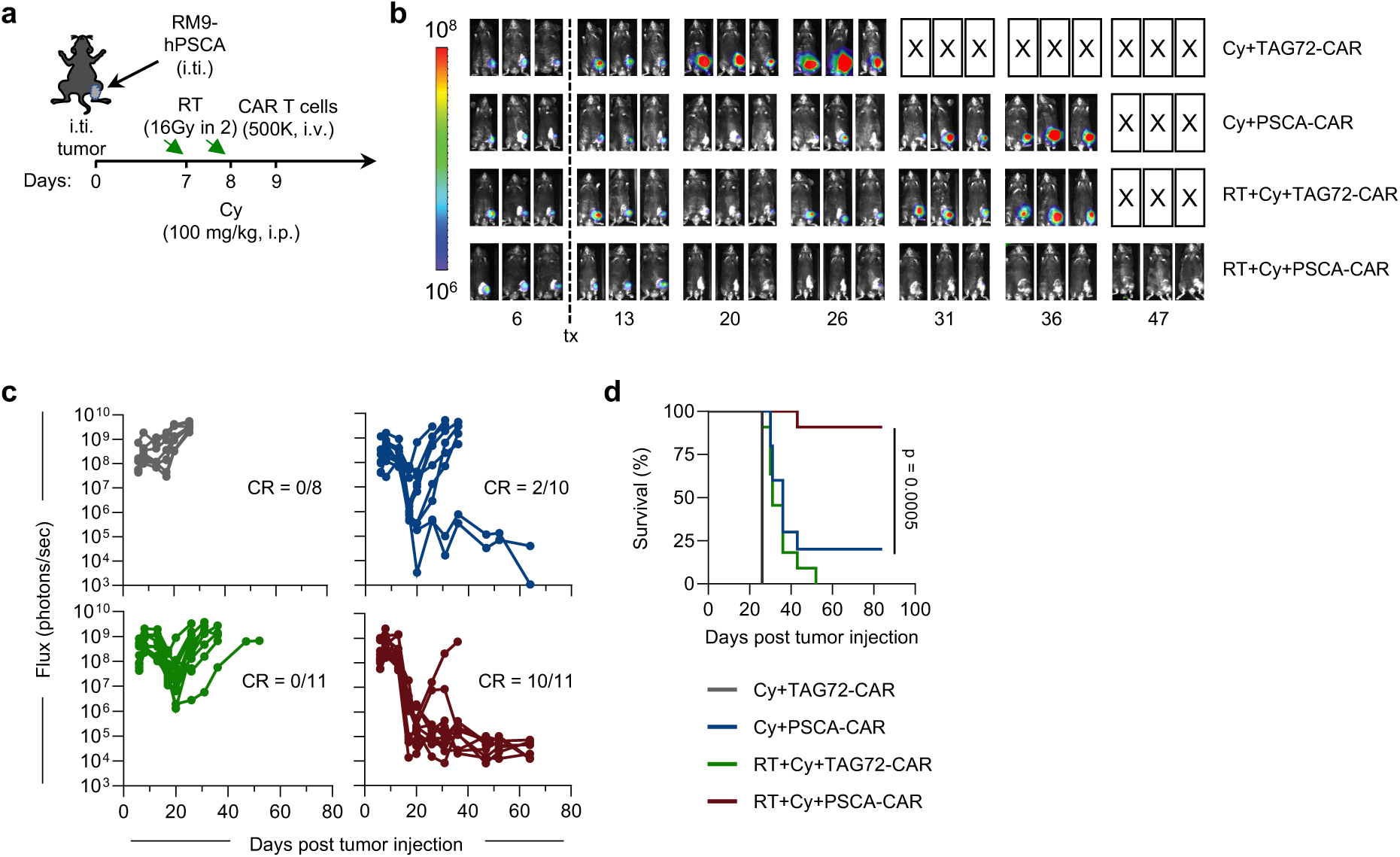
The combination of focal RT, Cy lymphodepletion, and PSCA-CAR T cells improves survival against bone-metastatic RM9-hPSCA prostate tumors. **a**, Illustration of RM9-hPSCA intratibial (i.ti.) tumor engraftment and treatment with focal RT (16 Gy total in 2 fractions given 24 hours apart), Cy preconditioning (100 mg/kg, i.p. following the second RT dose), and 0.5 x 10^6^ T cells (TAG72-CAR or PSCA-CAR, i.v.) on the indicated days. **b**, Representative images of tumor bioluminescent flux at the indicated days post tumor injection in each treatment group. **c**, Quantification of tumor flux (photons/sec) as measured by bioluminescent imaging in each replicate mouse at indicated days post tumor injection. Signals were quantified using a consistent region of interest (ROI) for each animal; CR, complete response. **d**, Kaplan-Meier survival plot for mice in each indicated group; n=8-11 mice per group, p-value compares RT+Cy+PSCA-CAR and Cy+PSCA-CAR using a log-rank (Mantel-Cox) test.

### Focal RT shows dose-dependent tumor growth inhibition and immunomodulatory effects on irradiated tumors, but with limited effects on non-irradiated distal tumors

While we confirmed that focal RT stimulates immune landscape changes in the local TME, the impact of RT on the TME of a non-irradiated distal tumor was unknown. Some case reports for renal cell carcinoma and melanoma have described how focal RT given to a primary tumor leads to the regression of tumors at other sites outside of the irradiated field; however, it is unknown whether this phenomenon would occur in immunologically cold prostate tumors[19–23]. To address this, we engrafted two s.c. RM9-hPSCA mouse prostate tumors at different sites to model multifocal disease. One tumor site was untreated while the other site was irradiated with the following doses and corresponding biological equivalent dose (BED): 1) a low dose of 8 Gy total in 2 fractions with BED of 11.2 Gy_10;_ 2) a medium dose of 16 Gy total in 2 fractions with BED of 28.8 Gy_10;_ 3) a non-fractionated medium dose of 13 Gy total in 1 fraction with a BED of 29.9 Gy_10_; and 4) a high ablative dose of 32 Gy total in 2 fractions with BED of 83.2 Gy_10_ (**Figure 3a**). Tumor volume measurements showed that RT led to a dose-dependent growth inhibition of the irradiated tumor with limited effects on growth of the non-irradiated tumor (**Figure 3b**). There were also no differences in survival between radiation doses due to the unaffected growth of the non-irradiated tumor (**Figure 3c**). These results indicate that RT given to one tumor has limited cytotoxic effects on non-irradiated tumors in this model.

**Figure 3.**
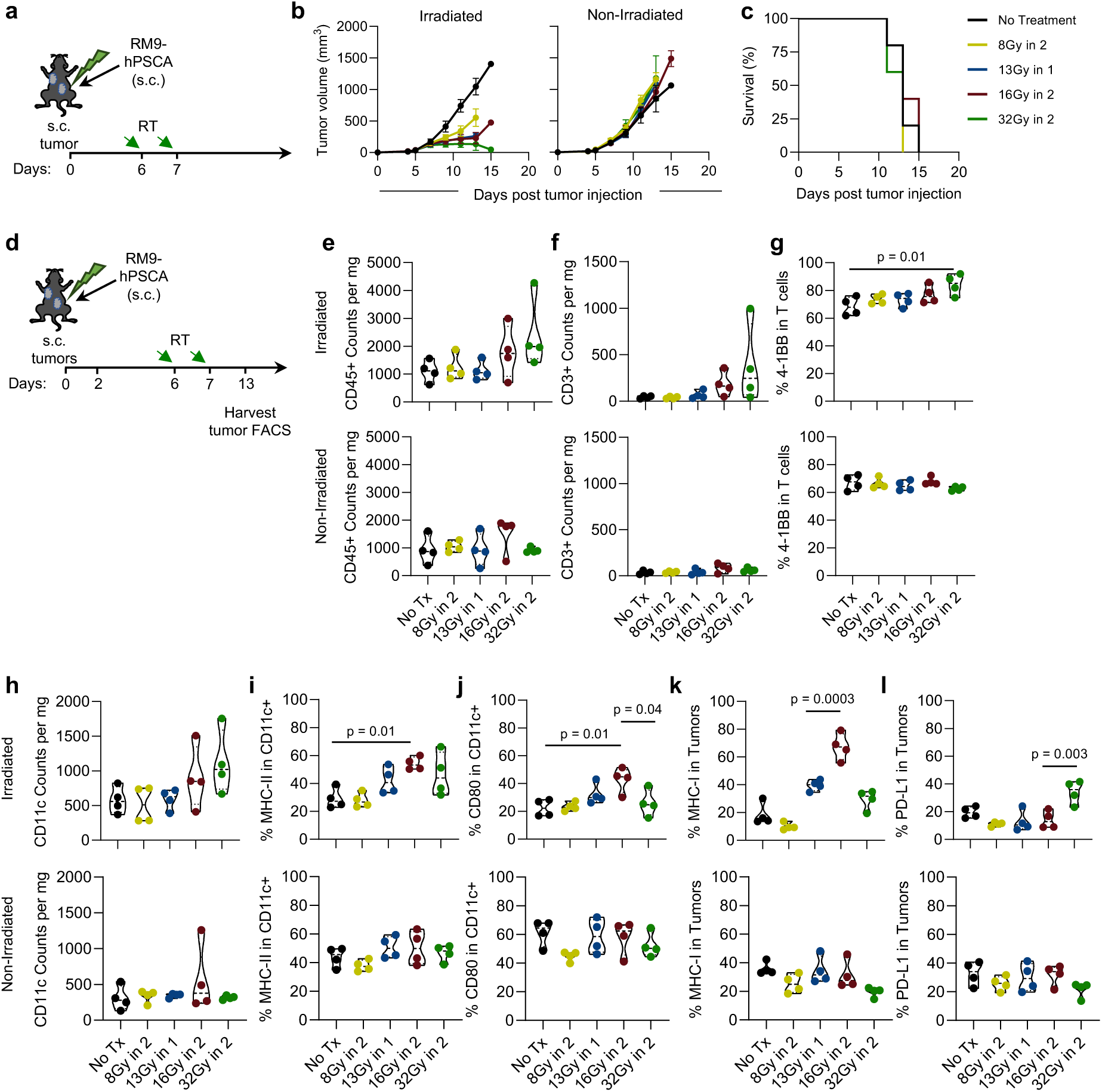
Focal RT induces dose-dependent growth inhibition and modulation of the TME in the irradiated tumors, but with limited effects on the non-irradiated tumors. a, Illustration of two RM9-hPSCA s.c. tumor engraftments and treatment with different doses of focal RT (8 Gy total in 2 fractions, 13 Gy total in 1 fraction, 16 Gy total in 2 fractions, and 32 Gy total in 2 fractions) on the indicated days post tumor injection. Focal RT given in 1 fraction was given 6 days post tumor injection and focal RT given in 2 fractions was given 6 and 7 days post tumor injection, 24 hours apart. **b**, Average tumor volumes (mm^3^) of the irradiated and non-irradiated tumors presented as mean ± SEM at indicated days post tumor injection for treatment groups. **c**, Kaplan-Meier survival plot for mice in each indicated group; n=5 mice per group. **d**, Illustration of two RM9-hPSCA s.c. tumors engrafted two days apart and treatment with different doses of focal RT on the indicated days following the first tumor injection, followed by tumor harvest and flow cytometric analysis on day 13 post tumor injection. **e-l**, Flow cytometry analysis of irradiated and non-irradiated tumors harvested on day 13 post tumor injection; n=4 mice per group: counts per mg of CD45+ immune cells (**e**), counts per mg of CD3+ T cells (**f**), percentage of 4-1BB+ activated T cells (**g**), counts per mg of CD11c+ dendritic cells (**h**), percentage of MHC-II+ dendritic cells (**i**), percentage of CD80+ dendritic cells (**j**), percentage of MHC-I+ tumor cells (**k**), and percentage of PD-L1+ tumors (**l**). In truncated violin plots (e-l), the curve of the plots extends to the minimum and maximum values, the center dashed line denotes the median value, and dotted lines represent quartiles. Unless otherwise indicated, p-values for pairwise comparisons were generated using the ordinary one-way ANOVA using Tukey’s multiple comparisons test.

We next examined the effects of different RT doses on the local immune landscape as well as at the nonirradiated site. In this study, two s.c. RM9-hPSCA tumors were staggered in engraftment by two days to model metachronous disease. The tumor engrafted earlier was irradiated with the same doses described in the previous experiment (**Figure 3d**). Both the irradiated and nonirradiated tumors were then harvested six days after RT treatment, or day 13 post engraftment of the first tumor, for flow cytometric analysis. The irradiated tumors showed greater infiltration of CD45+ immune cells, along with greater CD3+ T cell infiltration and 4-1BB activation with higher doses of RT (**Figure 3e-g, upper**). Higher RT doses also led to greater infiltration of CD11c+ DCs (**Figure 3h, upper**). In the non-irradiated tumor, however, RT showed limited changes in immune cell infiltration (**Figure 3e-g, lower**). Interestingly, the medium fractionated RT dose of 16 Gy total in 2 fractions significantly increased expression of the antigen presentation molecule MHC-II and the activation marker CD80 on CD11c+ DCs compared to no RT, lower doses, or the highest dose of 32 Gy in 2 fractions (**Figure 3i-j, upper**). This dose of RT also upregulated the expression of MHC-I on tumor cells compared to the other doses, while also not upregulating the checkpoint molecule PD-L1 (**Figure 3k-l, upper**). There were minimal immunological changes with any of the RT doses in the non-irradiated tumor. Considering that antigen presentation is an important aspect of activating the endogenous immune system, this data suggests that 16 Gy total in 2 fractions had the greatest beneficial and immune-stimulating effects on the local TME and was the dose used for subsequent combinatorial experiments with CAR T cells.

### The combination of focal RT, Cy lymphodepletion, and PSCA-CAR T cell treatment shows improved survival against a multifocal RM9-hPSCA mouse prostate tumor model

Although focal RT alone may not lead to regression in non-irradiated lesions, both preclinical and clinical studies have found that the combination of focal RT with immunotherapies leads to systemic immune responses against non-irradiated tumors[14, 24–29]. To test this, we engrafted two s.c. RM9-hPSCA tumors, irradiated one of the tumors with 16Gy in 2 fractions, and treated mice with Cy and CAR T cells (**Figure 4a**). Non-targeting Cy+TAG72-CAR T cells and RT+Cy+TAG72-CAR T cell treatments transiently inhibited the growth of both tumors (**Figure 4b**). This suggests that irradiation of the first tumor with Cy preconditioning and without concurrent tumor-specific T cell therapy failed to control the growth of the non-irradiated tumor. The Cy+PSCA-CAR T cell treatments affected the growth of both tumors similarly, as expected, curing 4 of 26 mice (**Figure 4b-c**). The triple combination of RT+Cy+PSCA-CAR T cells was able to completely cure 23 out of 26 of the irradiated tumors, which reflects the potent antitumor response seen in the single tumor models. This combination treatment also cured more of the non-irradiated tumors compared to the other treatments, leading to the greatest survival benefit, though it did not reach statistical significance (*p* = 0.075) compared to the Cy+PSCA-CAR T cell treatment (**Figure 4c**). Minimal toxicities by body weight were observed with any treatment group (**Figure 4d**). Together, these studies suggest that the RT+Cy+PSCA-CAR T cell combination improves survival in a multifocal prostate cancer model.

**Figure 4.**
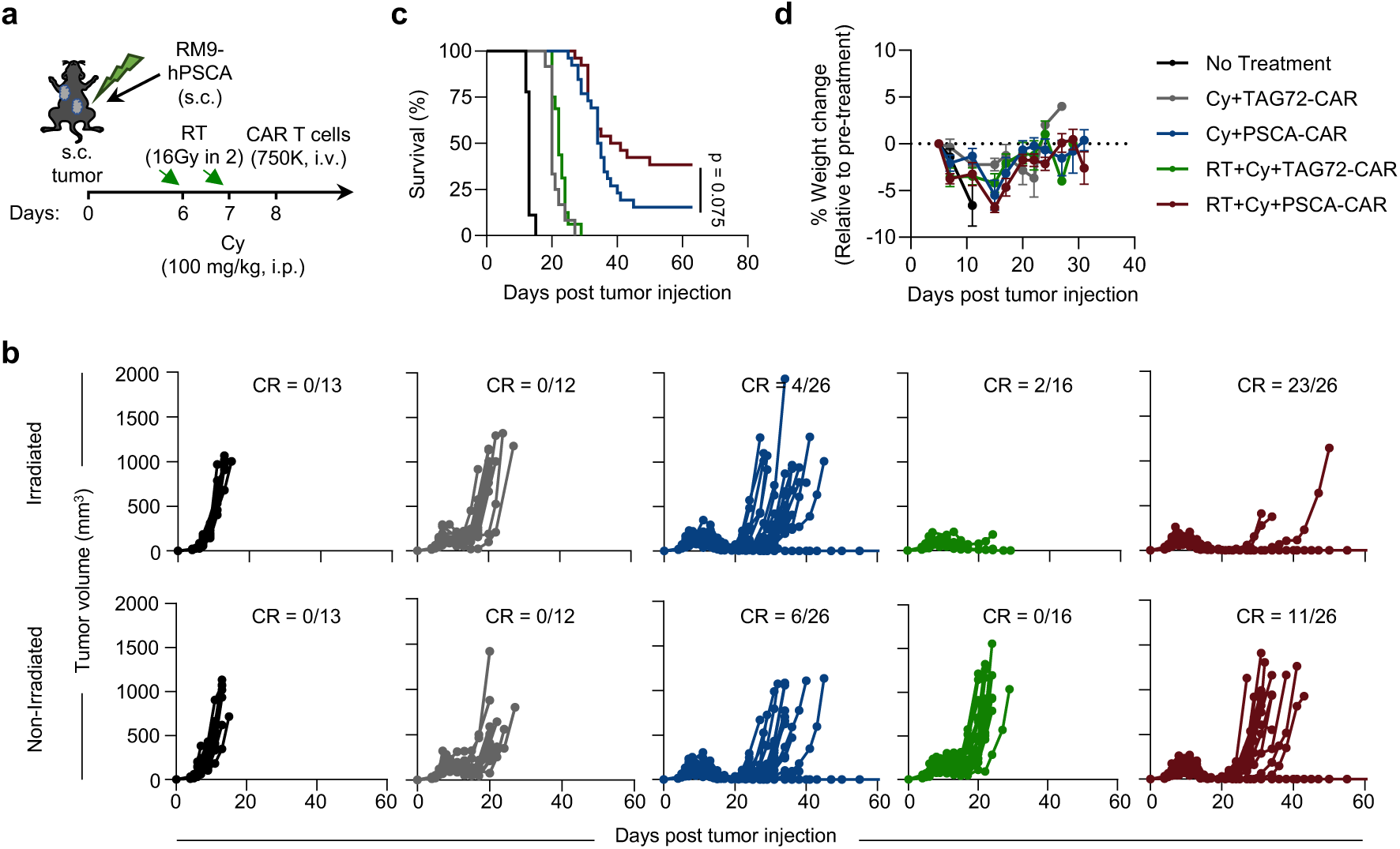
The combination of focal RT, Cy lymphodepletion, and PSCA-CAR T cells improves therapeutic responses against a multifocal RM9-hPSCA mouse prostate model. **a**, Illustration of two RM9-hPSCA s.c. tumor engraftments and treatment with focal RT (16 Gy total in 2 fractions given 24 hours apart), Cy preconditioning (100 mg/kg, i.p. following the second RT dose), and 0.75 x 10^6^ T cells (TAG72-CAR or PSCA-CAR, i.v.) on the indicated days. **b**, Tumor volume (mm^3^) measurements of each replicate at indicated days post tumor injection for treatment groups in the irradiated and non-irradiated tumor; CR, complete response. **c**, Kaplan-Meier survival plot for mice in each indicated group; n=13-26 mice per group, p-value compares RT+Cy+PSCA-CAR and Cy+PSCA-CAR using a log-rank (Mantel-Cox) test. **d**, Percentage body weight change of indicated groups relative to pre-treatment weight; n=5-12 mice per group, presented as mean ± SEM.

### The combination of focal RT, Cy lymphodepletion, and PSCA-CAR T cell treatment enhances antigen presentation in the TME and stimulates endogenous immunity in the tumor-draining lymph nodes

To better understand the effects of combining RT, Cy, and PSCA-CAR T cell therapy on local immune landscape changes of irradiated and non-irradiated tumors in our multifocal disease model, we harvested tumors four days after CAR T cell treatment (six days post-RT) and processed them for flow cytometric analysis. We observed approximately two-fold higher CD8/CD4 T cell ratios in the irradiated tumors of mice treated with RT+Cy+PSCA-CAR T cells compared to the No Treatment tumors while also trending higher than the other treatment groups (**Figure 5a**), suggesting that RT enhances the infiltration of effector CD8+ T cells to the irradiated tumor of RT+Cy+PSCA-CAR T cell-treated mice. The triple combination treatment also led to higher levels of the T cell activation marker 4-1BB, which could also contribute to greater cytotoxic effects at the irradiated tumor (**Figure 5b**). We did not observe significant differences in T cells between treatment groups in the non-irradiated tumor. We then compared the myeloid cell populations between treatment groups. In the irradiated tumor, RT+Cy+PSCA-CAR T cell treatment led to the highest frequency of MHC-II expression on CD11b+ myeloid cells compared to other treatment groups (**Figure 5c**). In the non-irradiated tumor, the mice that received PSCA-CAR T cell treatment showed similar upregulation of MHC-II molecules, which was higher than in other groups (**Figure 5c**). PD-L1 expression on CD11b+ myeloid cells was also significantly higher in the irradiated tumor of mice that received RT+Cy+PSCA-CAR T cells (**Figure 5d**).

**Figure 5.**
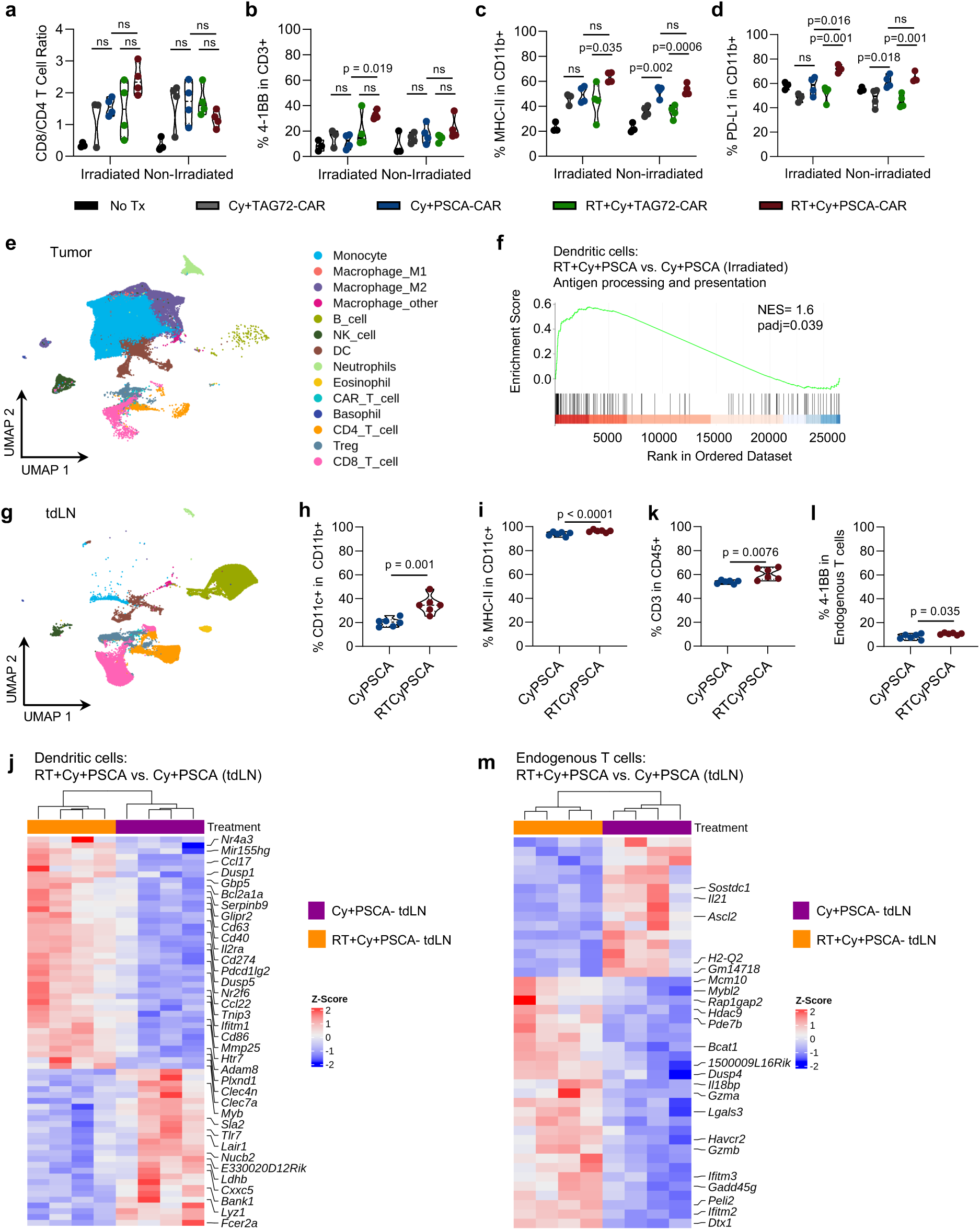
The combination of focal RT, Cy lymphodepletion, and PSCA-CAR T cells leads to immune changes in the irradiated TME and tdLN. **a-d**, Flow cytometry analysis of irradiated and non-irradiated tumors harvested 4 days post CAR T cell treatment; n=3-4 mice per group: CD8/CD4 T cell ratio (**a**), percentage of 4-1BB+ activated T cells (**b**), percentage of MHC-II+ CD11b+ myeloid cells (**c**), percentage of PD-L1+ myeloid cells (**d**). P-values for pairwise comparisons were generated using the ordinary one-way ANOVA using Tukey’s multiple comparisons test; ns, not significant. **e**, Uniform Manifold Approximation and Projection (UMAP) of immune cells within tumors harvested 5 days post CAR T cell treatment from Cy+PSCA-CAR T cell and RT+Cy+PSCA-CAR T cell-treated mice. **f**, Running enrichment score plot of antigen processing and presentation pathway in dendritic cells within tumors; padj, adjusted p-value. **g**, UMAP of immune cells within tumor-draining lymph nodes (tdLN) harvested 5 days post CAR T cell treatment from Cy+PSCA-CAR T cell and RT+Cy+PSCA-CAR T cell-treated mice. **h-i,k-l**, Flow cytometry analysis of tdLN’s harvested 5 days post CAR T cell treatment; n=6 mice per group: percentage of CD11c+ DCs (**h**), percentage of MHC-II+ CD11c+ DCs (**i**), percentage of CD3+ T cells (**k**), and percentage of 4-1BB+ activated endogenous T cells (**l**). Determination of p-values was generated using an unpaired Student’s t test. **j**, Heat map of differentially expressed genes within tdLN DCs. **m**, Heat map of differentially expressed genes within tdLN endogenous T cells. In truncated violin plots (a-d, h-i, k-l), the curve of the plots extends to the minimum and maximum values, the dashed center line denotes the median value, and the dotted lines represent the quartiles.

To understand the transcriptional changes generated by the RT+Cy+PSCA-CAR T cell and Cy+PSCA-CAR T cell treatments, we harvested tumors five days following CAR T cell treatment (seven days post-RT), processed tumors into single cell suspensions, enriched for tumor-infiltrating lymphocytes, and performed single-cell RNA-sequencing (scRNA-seq). Uniform Manifold Approximation and Projection (UMAP) plots were generated to visualize the immune cell populations (**Figure 5e**). RT+Cy+PSCA-CAR T cell treatment led to an upregulation of antigen processing and presentation pathways in multiple myeloid populations, including DCs, M1-like macrophages, and monocytes (**Figure 5f, Supplemental Figure 3a-b**). This is consistent with our flow cytometric analysis showing higher expression of MHC-II on the myeloid cells of RT+Cy+PSCA-CAR T cell-treated mice. Similar to the flow cytometric analyses of the nonirradiated tumor, transcriptional changes were undetected between the treatment groups.

Because the tumor-draining lymph nodes (tdLN) are critical sites for immune cell priming and immunotherapy responses, we also harvested the tdLN’s for scRNA-seq analysis and generated UMAPs for visualization (**Figure 5g**) [30–34]. We found that the RT+Cy+PSCA-CAR T cell treated mice had a higher frequency of CD11c+ dendritic cells in the tdLN that expressed higher levels of MHC-II molecules compared with Cy+PSCA-CAR T cell-treated mice (**Figure 5h-i**). Further analysis of the tdLN DCs revealed significantly higher expression of multiple immune-activating (*Cd40, Cd86, Ccl17, Il2ra*), co-inhibitory (*Cd274, Pdcd1lg2*), and interferon-stimulated gene signatures (*Ifitm1, Gbp5, Mir155hg*), indicating a strong immune priming response in the tdLNs (**Figure 5j**). We also analyzed the T cell population within the tdLN and found an elevated frequency of CD3+ T cells that also showed greater expression of the activation marker 4-1BB in the endogenous T cell population (**Figure 5k-l**). Using scRNA-seq analysis, the overall endogenous T cell population within the tdLNs of RT+Cy+PSCA-CAR T cell-treated mice showed upregulation of cytotoxic (*Gzma, Gzmb*), exhaustion (*Havcr2*), inflammatory interferon-stimulated (*Ifitm2, Ifitm3, Il18bp*), and proliferation-related genes (*Mcm10, Mybl2*), suggesting that T cells are primed for cytotoxicity, antigen-experienced, and expanding in the tdLNs (**Figure 5m**). Overall, these results show that the RT+Cy+PSCA-CAR T cell treatment leads to a more inflammatory and antigen-visible TME that stimulates antigen presenting cells and T cells in the lymphoid tissue, leading to potent antitumor responses in our models of prostate cancer.

## Discussion

In this study, we investigated the efficacy of PSCA-CAR T cells in combination with focal RT using immunocompetent mouse prostate cancer models. We showed that focal RT alone induced dose-dependent immune cell infiltration and high levels of antigen presentation in the irradiated mouse prostate TME. We observed that the combination of focal RT, Cy preconditioning, and PSCA-CAR T cells led to robust and durable antitumor responses and overall survival in mouse prostate cancer models, including two different subcutaneous models and one clinically relevant bone metastatic prostate cancer model. Given the potency of the combination treatment at a single irradiated tumor site and the immunogenicity of focal RT treatment, we hypothesized that this combination treatment with focal RT given to one site would also stimulate systemic immunity and promote immune responses against non-irradiated lesions. Consequently, we found that this combination of treatments showed trends of improving survival against a multifocal prostate cancer disease model without toxicity. We examined the immune landscape changes induced by these treatments and observed upregulation of antigen processing and presentation pathways by multiple myeloid subtypes in the irradiated tumor. Furthermore, the tdLN contained dendritic cells that were more activated and primed for T cell stimulation, as well as endogenous T cells that were also more proliferative and cytotoxic compared to non-irradiated mice. Altogether, these studies demonstrate the potency of focal RT along with Cy preconditioning and PSCA-CAR T cell therapy against prostate cancers.

Other preclinical studies have shown the benefits of using RT alone to locally irradiate the tumor prior to CAR T cell therapy. The combination of RT and CAR T cells without Cy lymphodepletion improved survival compared to CAR T cells alone in both immunocompromised and immunocompetent mouse models[14–16, 35–38]. Our studies, however, showed that RT and CAR T cell treatments failed to show curative responses. This difference could be due to the different radiosensitivities, tumor sizes at the time of treatment, and immunosuppressive TME of each cancer model. For example, some studies showed that irradiating one tumor led to growth inhibition of a non-irradiated tumor in B cell lymphoma and pancreatic ductal adenocarcinoma, suggesting that those tumors may be more radiosensitive[16, 24]. Our studies demonstrate that the mouse prostate cancer TME is particularly suppressive, such that RT alone is insufficient to improve CAR T cell therapy. These studies indicate that RT improves antitumor responses; however, Cy preconditioning is crucial for enhancing CAR T cell cytotoxicity against prostate cancers.

Minimal toxicities were observed in our preclinical models as determined by measuring body weight change. However, in our phase 1 clinical trial assessing the safety and bioactivity of PSCA-CAR T cells, dose-limiting toxicities in the form of cystitis were observed in patients that received a higher dose of Cy lymphodepletion, which was mitigated upon lowering the Cy dose[4]. Patients that received a higher Cy dose also showed trends of greater CAR T cell expansion in the peripheral blood, indicating that excessive CAR T cell proliferation may exacerbate bladder cystitis. In light of this and despite observing minimal toxicities in our mouse models, the greater proliferation of T cells observed in the tdLN of irradiated mice suggests that close and frequent monitoring of patients receiving this combination treatment in clinical trials may be warranted.

We acknowledge that our study tested only a limited number of doses and that there is a wider RT dose range and regimen that can be explored. However, the doses tested are clinically relevant and were within the range of doses tested in clinical trials that were found to be relatively safe with minimal toxicities[39–45]. Furthermore, 8 Gy per fraction is reported to have lower rates of toxicity while maintaining biochemical control rates similar to higher fractional doses and is often used in palliative treatments for bone metastases[42, 46]. Consequently, this treatment regimen can be more easily adapted in the clinic in combination with other treatments that may benefit from the immunostimulatory effect of that radiation dose, such as immunotherapies, leading to higher efficacy and improved responses.

We acknowledge that we did not compare the effects of different timing and sequences of the combination treatments. In this study, we assessed the sequence of focal RT, followed by Cy lymphodepletion and CAR T cell treatment. However, RT has been shown to recruit regulatory T cells (Tregs) to the tumor, so subsequent Cy treatment may help eliminate Tregs before they can negatively impact CAR T cell activity[47, 48]. We did not test the sequence of Cy and CAR T cells followed by RT. Although there is concern that irradiating the tumor after CAR T cell treatment may be cytotoxic to the CAR T cells that infiltrate the tumor early and reduce antitumor efficacy, one study combining EGFR-CAR T cells with RT did not find significant differences in survival between the sequence of CAR T cell and RT treatments[15]. Future research is needed to understand how the sequence and timing of treatments will affect the overall therapeutic response in prostate cancer models.

Although we observed a higher frequency of curative responses in both tumors of the RT+Cy+PSCA T cell treatment in our multifocal disease model, the antitumor efficacy was weaker at the non-irradiated secondary site, highlighting the superior antitumor potency of directly irradiating the tumor combined with CAR T cell therapy. Given this, multifocal RT in combination with systemic immunotherapy treatment may be necessary in oligometastatic disease settings to allow for a more robust antitumor response[49]. Other studies combining RT with checkpoint inhibitors have also found that high-dose irradiation to one tumor combined with low-dose radiation to other sites improves responses to checkpoint inhibitors[50–53]. Some metastatic sites may be difficult to irradiate locally, however; additional studies are needed to determine whether systemic radiopharmaceuticals, such as Lutetium-177 PSMA therapy or Radium-223, may yield more widespread TME changes that benefit CAR T cell therapy against prostate cancer. Altogether, the results of our study have been instrumental in the design of our ongoing phase 1b clinical trial evaluating the safety and bioactivity of PSCA-CAR T cells in mCRPC patients with or without metastasis-directed radiation therapy to multiple tumor sites (NCT05805371).

## Materials and Methods

### Cell Lines

The Ras/myc transformed prostate cancer line RM-9 was a kind gift from Timothy C. Thompson (MD Anderson Cancer Center). The mouse prostate cancer cell line TRAMP-C2 was a kind gift from Marcin Kortylewski (City of Hope National medical Center). Both RM-9 and TRAMP-C2 were cultured in Dulbecco’s Modified Eagles Medium (DMEM, Life Technologies) containing 10% fetal bovine serum (FBS, Hyclone), 25 mM HEPES (Irvine Scientific), and 2 mM L-Glutamine (Corning TM, complete DMEM).

### DNA constructs and tumor lentiviral transduction

Tumor cells were engineered to express firefly luciferase (ffluc) and the human prostate stem cell antigen (hPSCA), as described previously[3, 54]. Briefly, RM-9 was sequentially transduced with epHIV7 lentivirus carrying the ffluc gene under the control of the EF1α promoter and the epHIV7 lentivirus carrying the hPSCA gene, which was also under the control of the EF1α promoter. TRAMP-C2 was transduced with epHIV7 lentivirus carrying the hPSCA gene and underwent fluorescence-activated cell sorting (FACS) to generate cells that uniformly expressed high levels of hPSCA.

The PSCA-CAR and non-targeting TAG72-CAR constructs were previously described[3, 55]. Briefly, the PSCA-CAR consists of a murine anti-PSCA scFv (1G8) with murine CD8 hinge, murine CD8 transmembrane domain, murine 4-1BB costimulatory domain, and a murine CD3ζ cytoplasmic domain[56–58]. The CAR construct was separated from the truncated murine CD19 tag by a T2A ribosomal skip sequence and cloned into the pMYs retrovirus backbone (Cell BioLabs). The non-targeting TAG72-CAR construct was cloned into a similar pMYs retrovirus backbone but has a murine anti-TAG72 scFv (CC49)[59]. Retrovirus was generated using PlatE cells (Cell Biolabs) and FuGENE HD transfection reagent (Promega), as described previously[3, 60]. The viral supernatant was collected after 24h, 36h, and 48h, pooled, and frozen at -80°C until further use.

### Murine T cell isolation, transduction, and ex vivo expansion

Murine T cell activation and transduction were performed as described previously[3, 60]. Briefly, spleens were obtained from male heterozygous hPSCA-KI mice. Mouse T cell enrichment was performed using the EasySep mouse T cell isolation kits according to the manufacturer’s protocol (StemCell Technologies). Mouse T cells were activated with mouse CD3/CD28 Dynabeads (Invitrogen) and transduced with PSCA-CAR or TAG72-CAR retrovirus.

### In vivo animal studies

All animal experiments were performed under protocols approved by the City of Hope Institutional Animal Care and Use Committee. All studies used 6- to 8-week- old male heterozygous hPSCA-KI C57BL/6j mice, which were generated as described previously[3, 61]. The mice were co-housed in a maximum barrier, pathogen-and opportunist-free animal facility. Mice were euthanized by CO_2_ and cervical dislocation once the endpoint was reached or upon signs of distress, including apparent weight loss, impaired mobility, or evidence of being moribund.

For *in vivo* subcutaneous (s.c.) tumor studies, 1.0 x 10^6^ RM9-hPSCA/ffluc cells or TRAMPC2-hPSCA cells were resuspended in 100 µL Hanks’ Balanced Salt Solution without Ca^2+^, Mg^2+^, or phenol red (HBSS^-/-^) buffer and subcutaneously engrafted in 6- to 8-week- old male heterozygous hPSCA-KI mice. For experiments with two s.c. tumors, 1.0 x 10^6^ RM9-hPSCA/ffluc cells were resuspended in HBSS^-/-^ buffer and engrafted in different regions of the abdomen. For rechallenge studies, 0.05 x 10^6^ RM9-WT tumors or 1 x 10^6^ TRAMPC2-WT tumors were prepared in 100 µL HBSS^-/-^ buffer and engrafted s.c. into tumor-naïve or cured mice. Tumor growth was monitored at least twice per week via caliper measurements. Tumor volume was calculated using the formula V= Length x Width x Height. Caliper measurements continued until scheduled harvest or humane endpoints were reached.

Mice engrafted with one s.c. RM9-hPSCA/ffluc or TRAMPC2-hPSCA tumor received two doses of 8 Gy focal radiation, either 6 or 24 hours apart (16 Gy total in 2 fractions) once tumors reached 100 mm^3^. In mice engrafted with two s.c. tumors, only one tumor received radiation at the following doses: 8 Gy total in 2 fractions, 16 Gy total in 2 fractions, 13 Gy in 1 fraction, or 32 Gy total in 2 fractions. If two fractions were given, the mice were irradiated 24 hours apart over two days. If only one fraction was given, the mice were irradiated on the first day. Cyclophosphamide (Cy) (Sigma-Aldrich) was given intraperitoneally (i.p.) at 100 mg/kg following the second dose of radiation treatment. Mice were then treated with the indicated dose of PSCA-CAR and TAG72-CAR T cells via retro-orbital intravenous (r.o. i.v.) treatment 24 hours after Cy treatment.

For mice engrafted with intratibial (i.ti.) tumors, 2.5 x 10^4^ RM9- hPSCA/ffluc cells were resuspended in 30 µL HBSS^-/-^ buffer and engrafted into the tibia of 6- to 8-week-old male mice as previously described[54]. Tumor growth was monitored at least once per week via biophotonic flux imaging (LagoX, Spectral Instruments) and quantified using Aura In Vivo Imaging Software (Spectral Instruments Imaging). Radiation treatment was given on day 7 and 8 post-engraftment, followed by i.p. Cy treatment on day 8 post-engraftment following final radiation treatment, and r.o. i.v. treatments with 0.5 x 10^6^ PSCA-CAR or TAG72-CAR on day 9 post-engraftment. Flux imaging continued until humane endpoints were reached.

### Irradiation

The radiation treatment was delivered by using the X-Rad Small-Animal Radiation Therapy (SmART)+ Biological Irradiator (Precision X-Ray, North Branford, CT, USA), a 3D image-guided, focused X-ray irradiation system operating at 225 kV and 13 mA. Prior to irradiation, animals were anesthetized with inhaled isoflurane and oxygen and subjected to cone-beam computed tomography (CBCT) for treatment planning. The acquired CBCT images were used to define the radiation beam layout (size and positioning), ensuring precise alignment of the isocenter within the tumor while avoiding exposure to adjacent critical structures such as bone.

CT-guided radiation treatment planning was performed using SmART-Advanced Treatment Planning (ATP) (SmART Scientific Solutions B.V.), a Monte Carlo-based software specifically developed for small animal research. Radiation treatment was given using a 10 x 10 mm^2^ (1 cm^2^) square collimator using two laterally opposing parallel beams, with the dose normalized to the isocenter located at the tumor’s midpoint.

### Flow Cytometry

Flow cytometric processing and analyses were performed as previously described[3, 54, 55, 60]. Briefly, subcutaneous tumors and tissues were harvested from mice, minced, and enzymatically digested into single cell suspensions using Dnase I (Invitrogen) and Collagenase D (Invitrogen) or a Miltenyi mouse tumor digestion kit (Miltenyi Biotec) and resuspended in HBSS^-/-^ with 2% FBS. Samples were incubated with rat anti-mouse Fc Block (BD Biosciences, Cat: 553142) at room temperature for 15 minutes followed by incubation for 30 minutes at 4°C in the dark with antibodies conjugated with eFluor506, Brilliant Violet 510 (BV510), VioGreen, Brilliant Violet 570 (BV570), Brilliant Violet 605 (BV605), Brilliant Violet 650 (BV650), fluorescein isothiocyanate (FITC), phycoerythrin (PE), PE/Dazzle 594, PE-cyanine (PE-Cy5), peridinin chlorophyll protein complex (PerCP), Percp-eFluor710, PE-Cy7, allophycocyanin (APC), Alexa Fluor 647 (AF647), red 718 (R718), and APC-Cy7. Anti-mouse antibodies include: PD-1 (Invitrogen, Cat: 69-9985-80, Clone: J43), NK1.1 (BioLegend, Cat: 108733, Clone: PK136), CD11b (BioLegend, 64 Cat: 101237, Clone: M1/70), LAG3 (BioLegend, Cat: 125227, Clone: C9B7W), CD3 (BioLegend, Cat: 100204, Clone: 17A2), TIM3 (BioLegend, Cat: 119704, Clone: RMT3-23), CD45 (BioLegend, Cat: 103145, Clone: 30-F11), CD44 (Invitrogen, Cat: 103010, Clone: IM7), CD4 (Invitrogen, Cat: 46-0042-82, Clone: RM4-5), 4-1BB (Invitrogen, Cat: 25-1371-82, Clone: 17b5), CD62L (BioLegend, Cat: 104412, Clone: MEL-14), CD19 (BD Bioscences, Cat: 567063, Clone: 1D3), CD8a (BioLegend, Cat: 100714, Clone: 53-6.7), CD80 (BD Biosciences, Cat: 740130, Clone: 16-10A1), Ly6-C (BioLegend, Cat: 128029, Clone: HK1.4), MHCII (I-A/I-E) (BioLegend, Cat: 101237, Clone: M5/114.15.2), CD45 (BioLegend, Cat: 103108, Clone: 30-F11), CD163 (BioLegend, Cat: 155316, Clone: S15049I), CD11c (BioLegend, Cat: 117316, Clone: N418), F4/80 (BioLegend, Cat: 123127, Clone: BM8), CD103 (BioLegend, Cat: 121426, Clone: 2E7), PD-L1 (BioLegend, Cat: 124312, Clone: 10F.9G2), CD3 (BD Biosciences, Cat: 567296, Clone: 17A2), Ly6-G (BioLegend, Cat: 127623, Clone: 1A8), CD86 (BioLegend, Cat: 159203, Clone: A17199A), MHC-II (I-A/I-E) (BioLegend, Cat: 107641, Clone: M5/114.15.2), CD45 (BD Bioscences, Cat: 561868, Clone: 30-F11), MHC-I (H-2Kb) (BD Bioscences, Cat: 567063, Clone: 1D3), goat anti-mouse (BD Biosciences, Cat: 550589), CXCR3 (BioLegend, Cat: 126506, Clone: CXCR3-173), and CD206 (BioLegend, Cat: 141706, Clone: C068C2). Unless specified otherwise, antibodies were diluted 1:100 for staining. Cell viability was determined using 4’, 6-diamidino-2-phenylindole (DAPI, Sigma, Cat: D8417). Flow cytometry was performed using the MACSQuant Analyzer 10 or the MACSQuant Analyzer 16 (Miltenyi Biotec). Data analysis was performed using FlowJo software (FlowJo LLC, v10.10).

### Single-cell RNA-sequencing and analysis

Mouse tumors and tumor-draining lymph nodes were processed into single cells as stated earlier. Tumor-infiltrating lymphocytes (TILs) were enriched from tumor samples using mouse CD4/CD8 (TIL) microbeads (Miltenyi Biotec) and pooled together with the flow-through. Single-cell gene expression libraries were generated using the Chromium GEM-X Single Cell ’5’ Gene Expression Kit v3 (10X Genomics, CA) according to the manufacturer’s instructions. Sequencing was performed using the Novaseq X platform (Illumina Inc., CA) with a sequencing depth of 30,000 reads per cell. Reads were aligned to the mm10 mouse reference genome, and expression matrices were generated using the Cell Ranger v9 software (10X Genomics). We added a segment of the CAR plasmid sequence to the reference genome and to the transcriptome annotations. Downstream analyses were performed using the R package Seurat v 5.3. Cells with more than 6,000 or fewer than 200 detected genes, and more than 20% mitochondrial UMIs were filtered out. The raw counts were normalized and log-transformed to use for downstream analyses. The highly variable features were identified using the FindVariableFeatures function to obtain 2,000 features per dataset. Next, a principal component analysis (PCA) was performed, and Uniform Manifold Approximation and Projection (UMAP) was used to perform dimensionality reduction using the first 30 principal components (PCs). Cell type annotations were made using canonical marker genes and the ScType package. Differential gene expression analyses and shrinkage of log2 fold changes were made on pseudo-bulk expression data from cells clustered by cell type using the DESeq2 v. 1.46.0 package. Pathway analysis was performed using the clusterProfiler v. 4.14.6 R package. Heat maps were made using the ComplexHeatmap v. 2.25.3 package.

### Statistical analysis and reproducibility

Data are presented as mean ± standard error of the mean (SEM) unless stated otherwise. Statistical comparisons between multiple groups were performed using the ordinary one-way analysis of variance (ANOVA) using Tukey’s multiple comparisons test to calculate the p-value unless otherwise stated. Statistical comparisons between two groups were performed using the unpaired Student’s t-test unless otherwise stated.

## Acknowledgements

We thank the staff of the following cores at the Beckman Research Institute at City of Hope (COH) Comprehensive Cancer Center: Animal Facility, Radiation Research Services, Pathology, Small Animal Imaging, and Integrative Genomics, and the Keck School of Medicine of the University of Southern California (USC) and the Norris Comprehensive Cancer Center Genomics Core for excellent technical assistance. Work performed was supported by the National Cancer Institute of the National Institutes of Health under grant numbers P30CA014089 (USC) and P30CA033572 (COH). Research reported in this publication was also supported by the National Cancer Institute of the National Institutes of Health under grant number F31CA278484 (PI: Young) and the Prostate Cancer Foundation (PIs: Priceman, Forman). The content is solely the responsibility of the authors and does not necessarily represent the official views of the National Institutes of Health.

## Author contributions

S.J.P., J.P.M., and C.A.Y. provided conception and construction of the study. S.J.P., C.A.Y., J.L., Y.R., R.R., L.C., J.P.M., designed the experimental procedures, data analysis, and/or interpretation. C.A.Y., J.L., Y.R., R.R., H.H., L.C., J.P.M., H.G., D.Z., A.M.H.A. performed experiments. C.A.Y. and S.J.P. wrote the manuscript. C.M., Y.Y., A.K.P., S.H., T.B.D., Y.R.L., and S.J.F. were consulted on methodologies and/or assisted in manuscript writing and editing. S.J.P. supervised the study. All authors reviewed the manuscript.

## Competing interests

S.J.P. is a scientific advisor and/or receives royalties from Imugene Ltd, Adicet Bio, Port Therapeutics, and Celularity. S.J.P. and S.J.F. are listed as co-inventors on a patent on the development of PSCA-targeted CAR-modified T cells for the treatment of PSCA-positive tumors, owned by City of Hope. All other authors declare that they have no competing interests.

