## Supplemental Figures for "Focal radiotherapy improves CAR T cell therapy targeting prostate cancer"

Figure S1

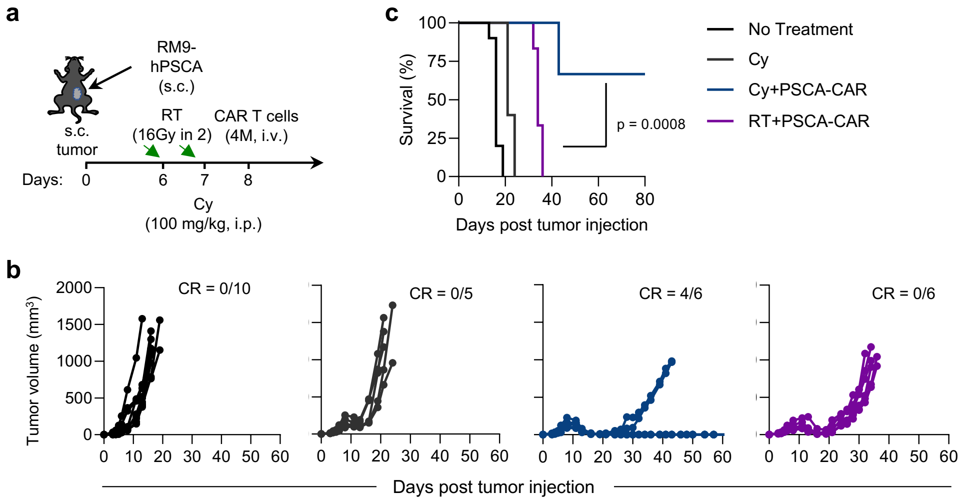


**Figure S1. Cy lymphodepletion is critical for curative responses with PSCA-CAR T cells against prostate cancer. a**, Illustration of RM9-hPSCA s.c. tumor engraftment and treatment with focal RT (16 Gy total in 2 fractions), Cy pre-conditioning (100 mg/kg, i.p.), and 4 x $10^6$ PSCA-CAR T cells (i.v.) at the indicated days. **b**, Tumor volume ($mm^3$) measurements of each replicate at indicated days post tumor injection for treatment groups; CR, complete response. **c**, Kaplan-Meier survival plot for mice in each indicated group; n≥5 mice per group, p-value compares RT+PSCA-CAR and Cy+PSCA-CAR using a log-rank (Mantel-Cox) test.

Figure S2

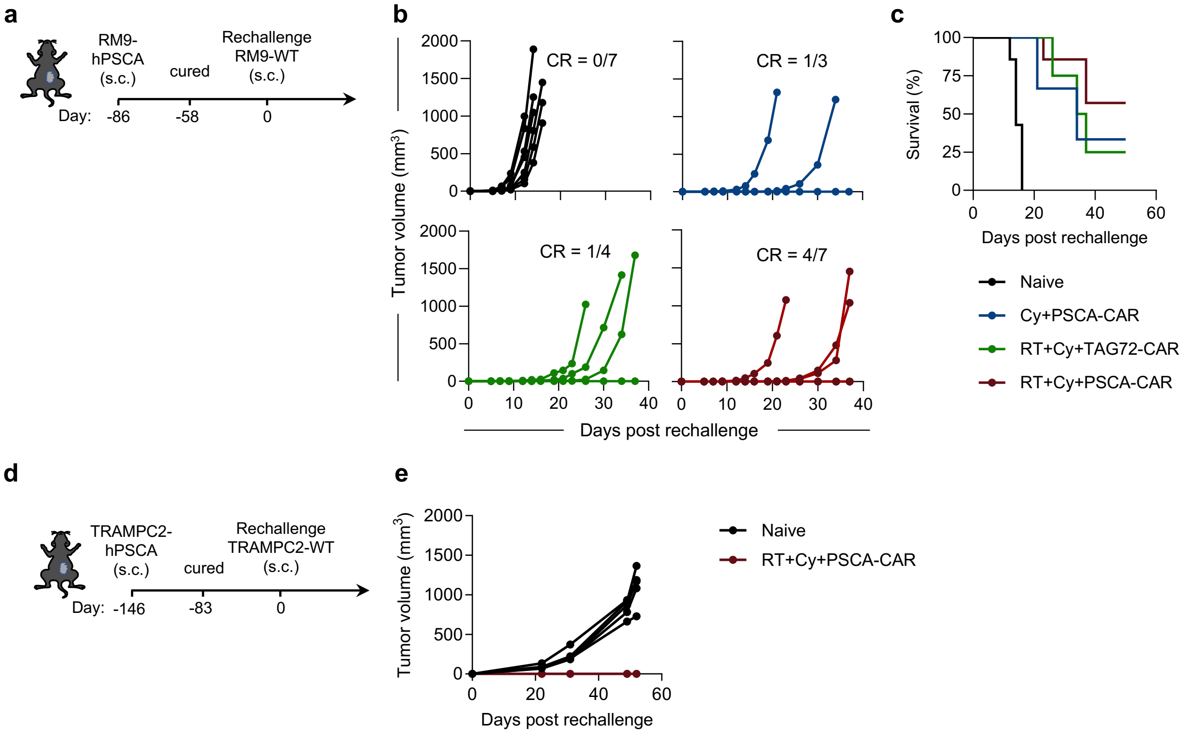


**Figure S2. The combination of focal RT, Cy lymphodepletion, and PSCA-CAR T cells induces a protective immune response.** **a**, Illustration of rechallenge with s.c. RM9-WT tumors 58 days following completely cured response. **b**, Tumor volume ($mm^3$) measurements of each replicate at indicated days post tumor injection for treatment groups; CR, complete rejection. **c**, Kaplan-Meier survival analysis for mice in each indicated group; n≥3 mice per group. **d**, Illustration of rechallenge with s.c. TRAMPC2-WT tumors 83 days following completely cured response. **e**, Tumor volume ($mm^3$) measurements of each replicate at indicated days post tumor injection for treatment groups; n=5 for Naïve mice and n=1 for RT+C+PSCA-CAR rechallenge.

Figure S3

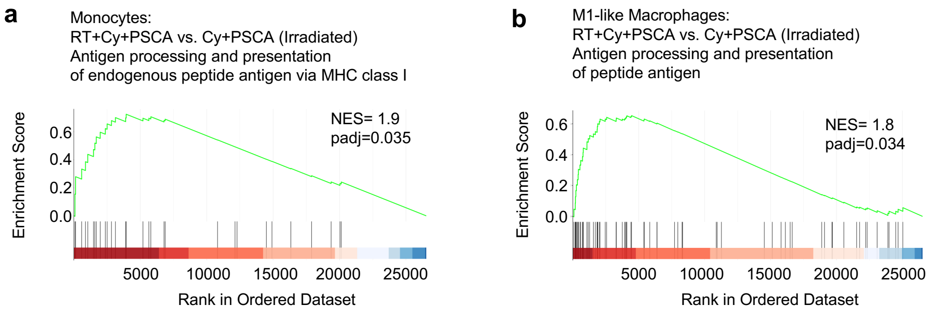


**Figure S3. Antigen processing and presentation pathways are upregulated in the monocytes and M1-like macrophages of irradiated tumors treated with focal RT, Cy lymphodepletion, and PSCA-CAR T cells. a-b,**, Running enrichment score plot of antigen processing and presentation of endogenous peptide antigen via MHC class I pathway in monocytes (**a**) and antigen processing and presentation of endogenous peptide antigen pathway in M1-like macrophages (**b**) within the irradiated tumors comparing RT+Cy+PSCA-CAR with Cy+PSCA-CAR; padj, adjusted p-value.
